# Comprehensive database of secondary metabolites from cyanobacteria

**DOI:** 10.1101/2020.04.16.038703

**Authors:** Martin R. Jones, Ernani Pinto, Mariana A. Torres, Fabiane Dörr, Hanna Mazur-Marzec, Karolina Szubert, Luciana Tartaglione, Carmela Dell’Aversano, Christopher O. Miles, Daniel G. Beach, Pearse McCarron, Kaarina Sivonen, David P. Fewer, Jouni Jokela, Elisabeth M.-L. Janssen

**Affiliations:** Department of Environmental Chemistry, Swiss Federal Institute of Aquatic Science and Technology (Eawag), 8600 Duebendorf, Switzerland; Centre for Nuclear Energy in Agriculture, University of São Paulo, CEP 13418-260 Piracicaba, SP, Brazil; School of Pharmaceutical Sciences, University of São Paulo, CEP 05508-900, São Paulo - SP, Brazil; Division of Marine Biotechnology, University of Gdansk, Al. Marszałka Piłsudskiego 46, 81-378, Gdynia, Poland; Department of Pharmacy, School of Medicine and Surgery, University of Napoli Federico II, Via D. Montesano 49, 80131 Napoli, Italy; Biotoxin Metrology, National Research Council Canada, 1411 Oxford Street, Halifax, Nova Scotia, B3H 3Z1, Canada; Department of Microbiology, University of Helsinki, 00014 Helsinki, Finland

## Abstract

Cyanobacteria form harmful mass blooms in freshwater and marine environments around the world. A range of secondary metabolites has been identified from cultures of cyanobacteria and biomass collected from cyanobacterial bloom events. A comprehensive database is necessary to correctly identify cyanobacterial metabolites and advance research on their abundance, persistence and toxicity in natural environments. We consolidated open access databases and manually curated missing information from the literature published between 1970 and March 2020. The result is the database CyanoMetDB, which includes more than 2000 entries based on more than 750 literature references. This effort has more than doubled the total number of entries with complete literature metadata and structural composition (SMILES codes) compared to publicly available databases to this date. Over the past decade, more than one hundred additional secondary metabolites have been identified yearly. We organized all entries into structural classes and conducted substructure searches of the provided SMILES codes. This approach demonstrated, for example, that 65% of the compounds carry at least one peptide bond, 57% are cyclic compounds, and 30% carry at least one halogen atom. Structural searches by SMILES code can be further specified to identify structural motifs that are relevant for analytical approaches, research on biosynthetic pathways, bioactivity-guided analysis, or to facilitate predictive science and modeling efforts on cyanobacterial metabolites. This database facilitates rapid identification of cyanobacterial metabolites from toxic blooms, research on the biosynthesis of cyanobacterial natural products, and the identification of novel natural products from cyanobacteria.

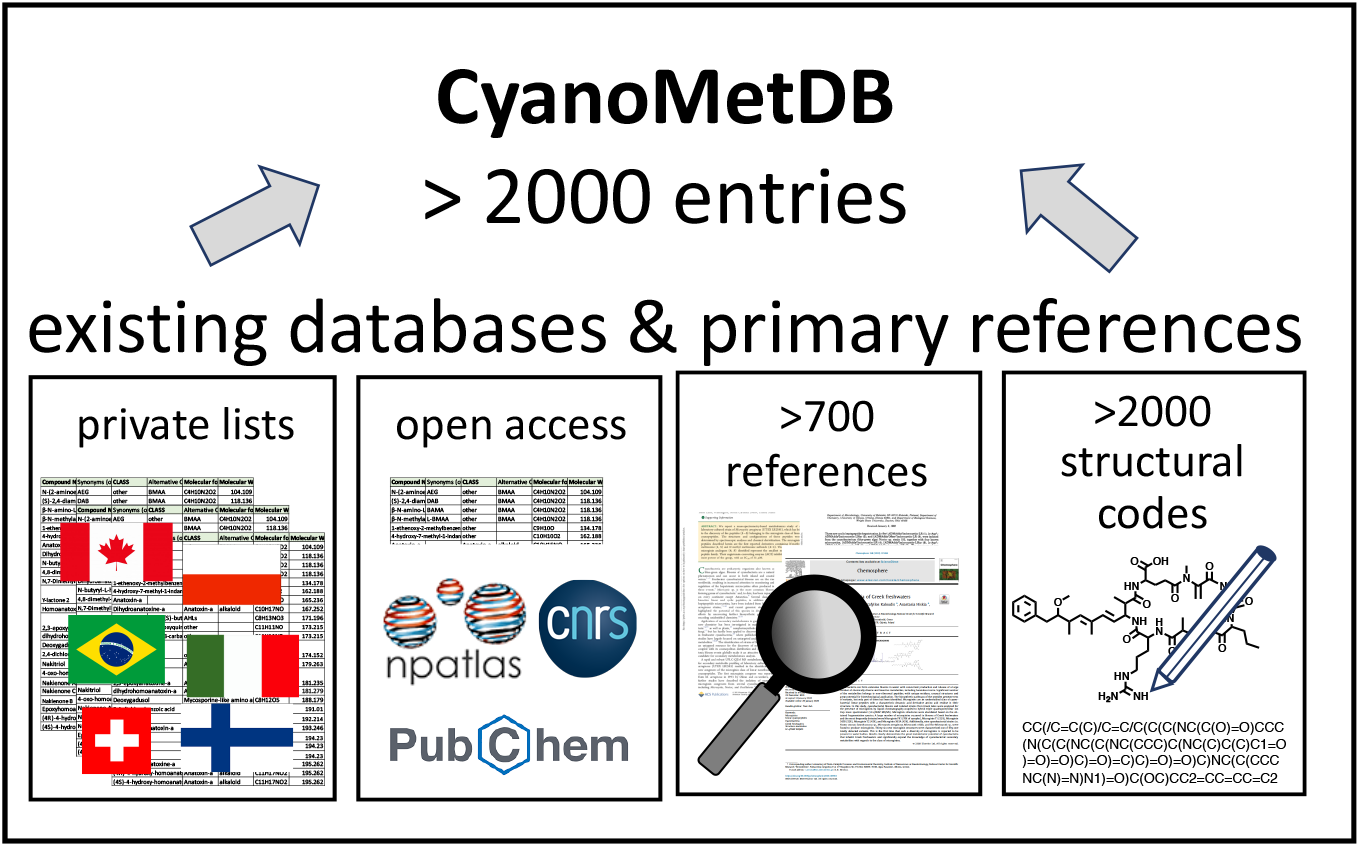

## Introduction

Around the globe, cyanobacteria inhabit fresh waters including drinking water reservoirs, brackish waters, and marine environments where they can proliferate to form harmful blooms. During these events, cyanobacteria can produce high concentrations of a diverse mixture of rather unique secondary metabolites. Various countries have put forward drinking water guidelines for one metabolite, microcystin-LR,^1^ for which the World Health Organization proposed a concentration threshold of 1 μg L^−1^ and the recently revised guidelines will also include the low molecular weight toxins anatoxin-a, saxitoxins and cylindrospermopsins.^2^ Currently, data on occurrence, fate, transformation processes, and toxicity of many other bioactive metabolites is lacking, and improved high-throughput analytical and effect-based methods are needed to overcome this information gap.

Research into analytical and toxicological methods relies on a comprehensive understanding of the structural information of cyanobacterial metabolites. One obstacle that contributes to this is the lack of a bioinformatics platform that the cyanobacteria researcher community collectively supports. While information from commercial databases of secondary metabolites are only accessible to paying customers (e.g., Antibase, MarinLit, The Dictionary of Natural Products), several open-access databases exist but are often limited regarding the number cyanobacterial metabolites or parameters listed (e.g., ALGALTOX List, NORINE database, Handbook of Marine Natural Products).^2–4^ Some key open access databases are listed in Table 1. The “Cyanomet mass” list by LeManach et al. (2019) contains 852 entries, of which 35 belong to the class of microcystins and nearly 500 compounds are listed with complete molecular formulae and literature reference but no further structural information is given.^3^ The Natural Product Atlas (2019) contains a similar number of entries for cyanobacterial metabolites, including microcystins, and also provides structural codes (e.g., SMILES code) and the stereochemistry is known for 768 entries (isomeric SMILES code).^4^ In 2017 the Handbook of Cyanobacterial Monitoring and Cyanotoxin Analysis published a list of 246 microcystins.^5^ Today, the most comprehensive list of microcystins and nodularins is curated by Miles et al. and was recently updated (2019) to include 279 microcystin variants with molecular formulae, references and a systematic naming system implying the structural compositions but no structural codes were provided.^6, 7^

**Table 1.** Publicly available suspect lists with total number of entries, number of microcystins, available molecular formula, primary reference for the structure elucidation and structural code (e.g., SMILES code).

| **Open Access List**<br>[reference] | **Entries** | **Microcystins** | **Mol. formula &**<br>**primary reference** | **Structural code**<br>**(e.g., SMILES code)** |
| --- | --- | --- | --- | --- |
| **CyanoMetMass<sup>3</sup>**<br>[LeManach et al. 2019] | 852 | 35 | 499 | 0 |
| **The Natural Product Atlas<sup>4</sup>**<br>[van Santen et al. 2019] | 887 | 34 | 887 | 887<br>(768 isomeric<br>SMILES codes) |
| **Microcystins_Miles<sup>6</sup>**<br>[Miles et al. 2019] | 279 | 279 | 279 | 0 |
| **CyanoMetDB**<br>[Jones et al. 2020] | **2049** | **310** | **2021** | **2031**<br>(1431 isomeric<br>SMILES codes) |

Here, we describe the compilation of information on secondary metabolites from cyanobacteria from these existing databases and literature curation into one database, CyanoMetDB. We first present the methodology for compiling the information, provide an overview of the content of the database, and summarize areas of research that can benefit from this database including suspect screening by mass spectrometry (MS) as well as analysis of biosynthetic variation, bioactivity, and environmental behaviour.

## Material and methods

### Parameter Selection

CyanoMetDB is a flat-file database (i.e., a single table) comprising the following core fields: compound identifier (key), compound name, compound class, molecular formula, molecular weight, monoisotopic mass, primary reference that elucidated the structure, detailed structural information as a simplified molecular input line entry system (SMILES codes) and other structural codes that serve as chemical identifiers for each compound (InChl, InChlKey, IUPAC names), whether an NMR platform was used for structure elucidation, and information about the sample used to elucidate the structure of the compound (genus/species/strain or *in situ* sample), see Table 2.^8^ Entries in the database correspond to compounds reported as cyanobacterial metabolites. In some instances, compounds reported in the database were first reported in non-cyanobacterial species, and have since been reported in one, or more, cyanobacterial species. In these cases, the reference for structure elucidation refers to the primary publication. The database lists secondary references where those compounds have later been reported for cyanobacteria as well. We indicate in the database which entries provide NMR spectroscopy data for confirming the planar structure. Other entries only offer tandem mass spectrometry (MS/MS) and potential mis-identification of some of these compounds is possible.

**Table 2.**
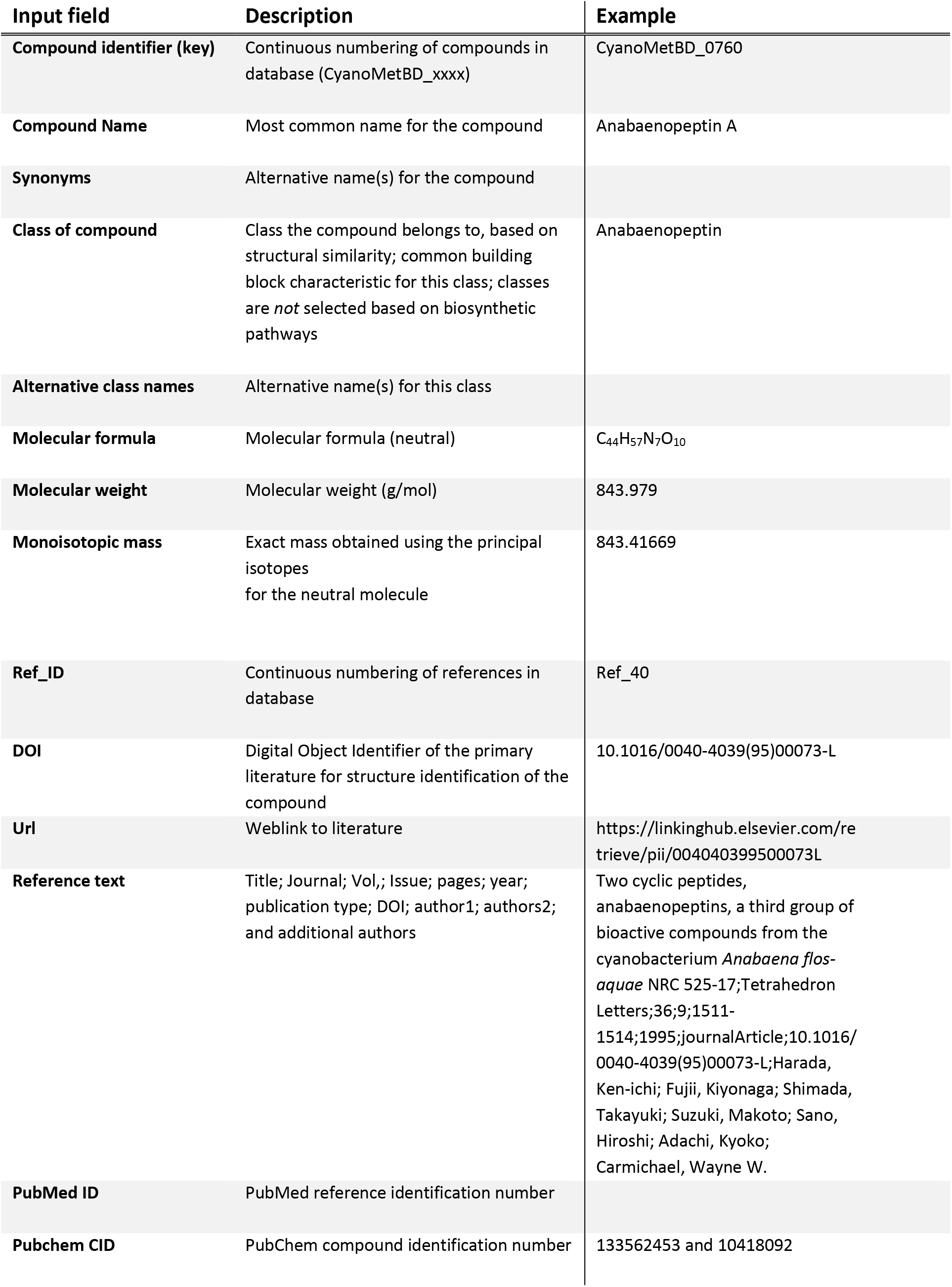
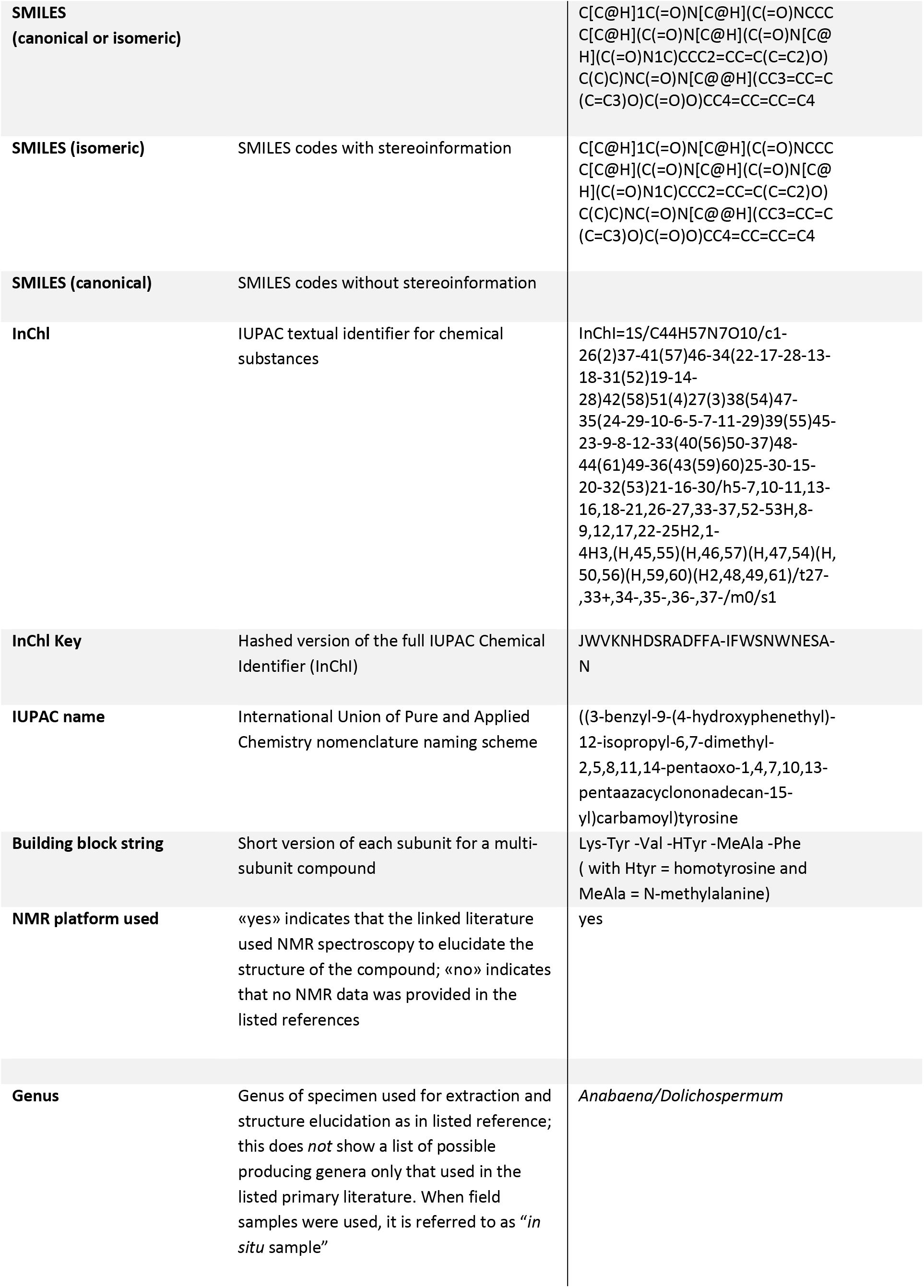
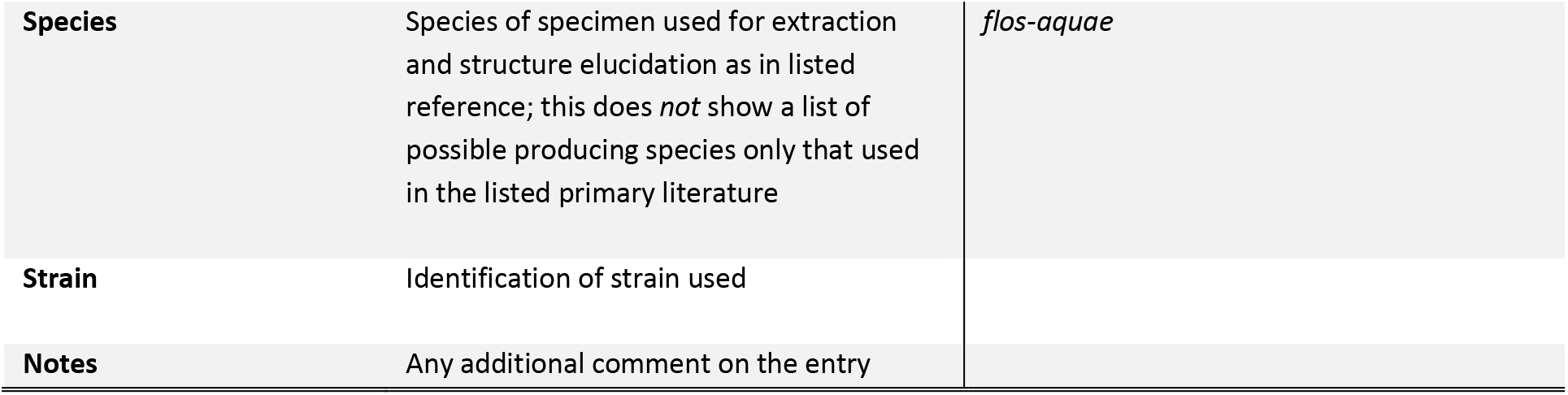
Description of the Individual Input Fields of Cyanobacterial Metabolite Database (CyanoMetDB).

| Input field | Description | Example |
| --- | --- | --- |
| **Compound identifier (key)** | Continuous numbering of compounds in database (CyanoMetBD_xxxx) | CyanoMetBD_0760 |
| **Compound Name** | Most common name for the compound | Anabaenopeptin A |
| **Synonyms** | Alternative name(s) for the compound |  |
| **Class of compound** | Class the compound belongs to, based on structural similarity; common building block characteristic for this class; classes are *not* selected based on biosynthetic pathways | Anabaenopeptin |
| **Alternative class names** | Alternative name(s) for this class |  |
| **Molecular formula** | Molecular formula (neutral) | C<sub>44</sub>H<sub>57</sub>N<sub>7</sub>O<sub>10</sub> |
| **Molecular weight** | Molecular weight (g/mol) | 843.979 |
| **Monoisotopic mass** | Exact mass obtained using the principal isotopes for the neutral molecule | 843.41669 |
| **Ref_ID** | Continuous numbering of references in database | Ref_40 |
| **DOI** | Digital Object Identifier of the primary literature for structure identification of the compound | 10.1016/0040-4039(95)00073-L |
| **Url** | Weblink to literature | https://linkinghub.elsevier.com/retrieve/pii/004040399500073L |
| **Reference text** | Title; Journal; Vol,; Issue; pages; year; publication type; DOI; author1; authors2; and additional authors | Two cyclic peptides, anabaenopeptins, a third group of bioactive compounds from the cyanobacterium *Anabaena flos-aquae* NRC 525-17;Tetrahedron Letters;36;9;1511-1514;1995;journalArticle;10.1016/0040-4039(95)00073-L;Harada, Ken-ichi; Fujii, Kiyonaga; Shimada, Takayuki; Suzuki, Makoto; Sano, Hiroshi; Adachi, Kyoko; Carmichael, Wayne W. |
| **PubMed ID** | PubMed reference identification number |  |
| **Pubchem CID** | PubChem compound identification number | 133562453 and 10418092 |
| **SMILES (canonical or isomeric)** |  | C[C@H]1C(=O)N[C@H](C(=O)NCCC[C@H](C(=O)N[C@H](C(=O)N[C@H](C(=O)N1C)CCC2=CC=C(C=C2)O)C(C)C)NC(=O)N[C@@H](CC3=CC=C(C=C3)O)C(=O)O)CC4=CC=CC=C4 |
| **SMILES (isomeric)** | SMILES codes with stereoinformation | C[C@H]1C(=O)N[C@H](C(=O)NCCC[C@H](C(=O)N[C@H](C(=O)N[C@H](C(=O)N1C)CCC2=CC=C(C=C2)O)C(C)C)NC(=O)N[C@@H](CC3=CC=C(C=C3)O)C(=O)O)CC4=CC=CC=C4 |
| **SMILES (canonical)** | SMILES codes without stereoinformation |  |
| **InChI** | IUPAC textual identifier for chemical substances | InChI=1S/C44H57N7O10/c1-26(2)37-41(57)46-34(22-17-28-13-18-31(52)19-14-28)42(58)51(4)27(3)38(54)47-35(24-29-10-6-5-7-11-29)39(55)45-23-9-8-12-33(40(56)50-37)48-44(61)49-36(43(59)60)25-30-15-20-32(53)21-16-30/h5-7,10-11,13-16,18-21,26-27,33-37,52-53H,8-9,12,17,22-25H2,1-4H3,(H,45,55)(H,46,57)(H,47,54)(H,50,56)(H,59,60)(H2,48,49,61)/t27-,33+,34-,35-,36-,37-/m0/s1 |
| **InChI Key** | Hashed version of the full IUPAC Chemical Identifier (InChI) | JWVKNHDSRADFFA-IFWSNWNESA-N |
| **IUPAC name** | International Union of Pure and Applied Chemistry nomenclature naming scheme | ((3-benzyl-9-(4-hydroxyphenethyl)-12-isopropyl-6,7-dimethyl-2,5,8,11,14-pentaoxo-1,4,7,10,13-pentaazacyclononadecan-15-yl)carbamoyl)tyrosine |
| **Building block string** | Short version of each subunit for a multi-subunit compound | Lys-Tyr -Val -HTyr -MeAla -Phe ( with HTyr = homotyrosine and MeAla = N-methylalanine) |
| **NMR platform used** | «yes» indicates that the linked literature used NMR spectroscopy to elucidate the structure of the compound; «no» indicates that no NMR data was provided in the listed references | yes |
| **Genus** | Genus of specimen used for extraction and structure elucidation as in listed reference; this does *not* show a list of possible producing genera only that used in the listed primary literature. When field samples were used, it is referred to as “*in situ* sample” | *Anabaena/Dolichospermum* |
| **Species** | Species of specimen used for extraction and structure elucidation as in listed reference; this does *not* show a list of possible producing species only that used in the listed primary literature | *flos-aquae* |
| **Strain** | Identification of strain used |  |
| **Notes** | Any additional comment on the entry |  |

We provide canonical SMILES codes representing the connectivity of each atom in a planar structure for all compounds and we aim to provide isomeric SMILES codes when stereochemical information was available from the literature. For some compounds, we list a second reference that improved or provided additional structural information following the initial structure elucidation. For some compounds, the “Note” section indicates where uncertainties for structure elucidation were observed. The stereochemistry of entries whose structures were established mainly through MS cannot be known with certainty, but where there was a reasonable basis for this, an assumed stereochemistry is included. For example, in the case of microcystins with fully established stereochemistries, position-2 and -4 have always been found to contain L-amino acids, with D-amino acids at position-1, 3 and -6. Similarly, the stereochemistry of di- and tetrahydrotyrosine in microcystins was assumed to be that established in other bacteria by Walsh et al.,^9, 10^ and this will if necessary be amended if found to be incorrect. We include some additional entries that are known metabolites, oxidation products, or (semi)synthetic and indicate them as such in the note section. However, we do not provide a comprehensive list of transformation products and synthetic compounds herein. The database also lists the material used to identify the compound from the primary literature as the genus and species of the cyanobacteria or whether it was a field sample. The database does not provide a comprehensive list of all known producing species.

Compounds are classified into major metabolite classes based on conserved molecular substructures as outlined in previous studies (Figure 1).^11^ For example: microcystins, heptapeptides with the characteristic Adda (i.e., 3*S*-amino-9*S*-methoxy-2*S*,6,8*S*-trimethyl-10-phenyl-deca-4*E*,6*E*-dienoic acid) derivatives; cyanopeptolins with a beta-lactone ring and the characteristic Ahp (3-amino-6-hydroxy-2-piperidone) moiety; anabaenopeptins with the ureido bond connecting the primary amine of lysine with the primary amine of the neighboring amino acid and lysine’s epsilon-amine with the carboxyl group of the C terminal amino acid to form a urea moiety and five membered peptide ring; aeruginosins with the Choi (2-carboxy-6-hydroxyoctahydroindole) and 3–4 substituents, and; microginins with the characteristic Ahda moiety (3-amino-2-hydroxydecanoic acid, or -octanoic acid) (Figure 1). We conducted substructure searches to verify the class assignments and identify other structural motifs (ChemAxon JChem Base software 20.4.0, 2020).

**Figure 1.**
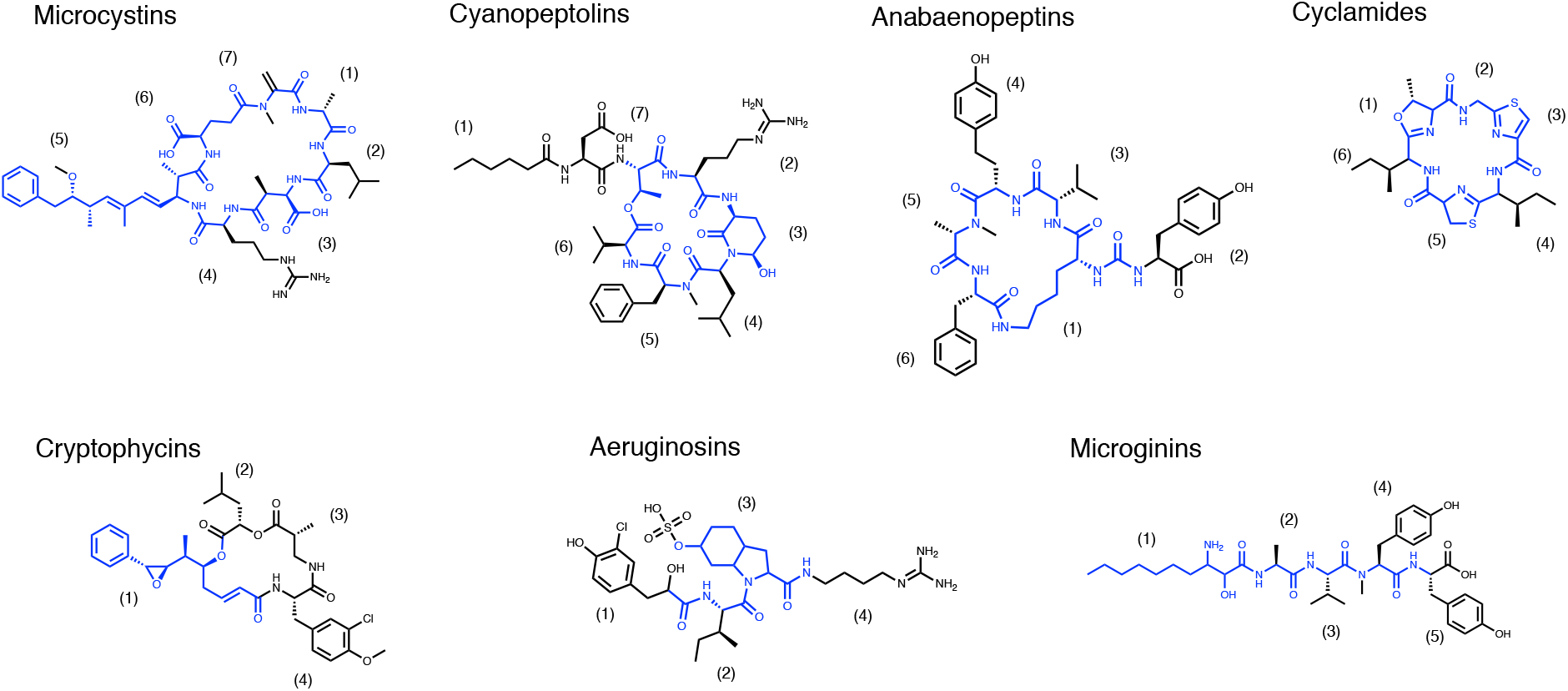
Representative compounds for seven cyanopeptide classes: microcystin-LR, cyanopeptolin A, anabaenopeptin A, aerucyclamide A, cryptophycin 1, aeruginosin 98A, and microginin 713. Core-structures that are shared among most variants in the respective class are marked in blue.

### Data sources and curation procedure

We manually verified and completed entries for the associated literature references using various bibliographic sources (e.g., existing open access databases, Sci-Finder, Pub-Med). From these references we extracted information of the sample type used therein (e.g., genus, species, field sample) and whether NMR spectroscopy was used for structure elucidation. Finally, with reference to the literature information, the molecules were manually drawn including known (or in some cases probable) stereochemistry and SMILES codes etc. were generated. In this way, some disagreements with the literature were identified and corrected. Entries extracted from PubChem were carefully checked to verify the structure from the primary literature. In case of discrepancies between structures reported in the primary literature reference and the one found in PubChem, we refer to the information derived from the primary literature reference herein. For selected compounds, remarks are included in the “Notes” field to highlight any shortcomings or relevant additional information, e.g., assumed stereochemistry of a compound. Information in Table 2 provides a detailed content description and illustrative example for each data field.

## Results and discussion

The database includes 2031 entries with complete literature and structural information. The earliest entry was published in 1970 for the cyanopeptolin micropeptin-996 followed by almost 100 additional compounds until 1990. Between 1990 and 2000, the number of reported cyanobacterial metabolites increased five-fold. The rapid increase during the 1990s was, in part, likely associated with the discovery that microcystins pose significant hepatotoxic risks to humans, which in turn lead to MC-LR being included in the World Health Organization’s water quality guidelines, prompting significant research on cyanobacteria.^2^ More than one thousand compounds were identified by the year 2010, and a further one thousand compounds have been reported in the subsequent decade (to March 2020). The historic data show that 50 to 130 compounds have been identified annually (Figure 2A). Increased availability of advanced analytical instrumentation has also contributed to the increased rate of discoveries in recent years (e.g., HRMS, high-field NMR spectroscopy with cryogenic probes).

**Figure 2.**
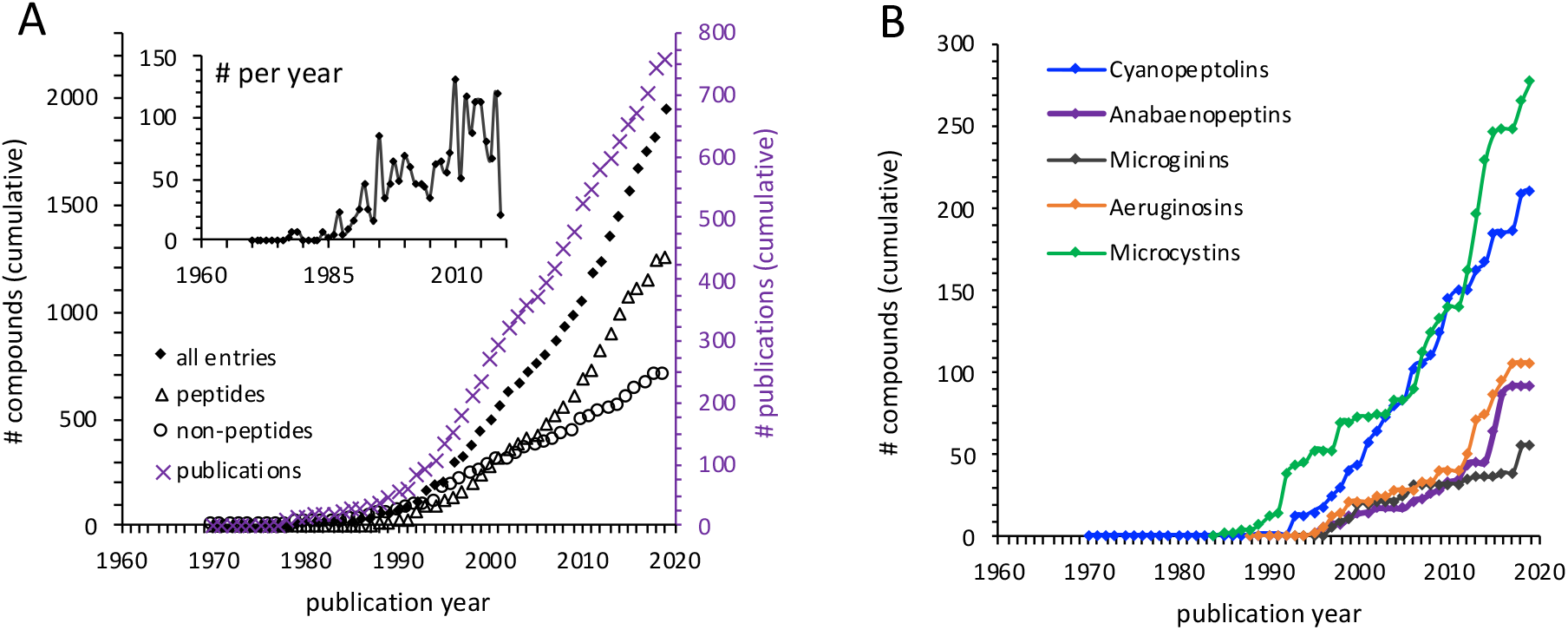
**(A)** Cumulative number of identified secondary metabolites from cyanobacteria (black diamonds); only peptide-based compounds (black open triangles); only non-peptide compounds (black open circles), and; number of publications (purple, secondary y-axis). Inset shows the number of compounds published each year (non-cumulative). **(B)** Cumulative numbers of major cyanopeptide classes of cyanopeptolins (blue); anabaenopeptins (purple); microginins (grey); aeruginosins (orange), and; microcystin (green) between 1970 and 2019.

### Publication trends

The discovery of cyanobacterial metabolites has not yet reached a plateau (Figure 2A). The question arises: How close we to identifying the majority of these cyanobacteria-specific metabolites? Also, how many of the newly described compounds are chemical variants of known classes and how many describe new classes? These are also overarching questions in the general domain of natural product research. Linginton et al. recently surveyed the chemical space of natural products and concluded that nowadays new discoveries mostly relate structurally to previously published compounds and that the “range of scaffolds readily accessible from nature is limited”.^12^ This does not mean that new discoveries are expected to be exhausted but rather that most new discoveries are likely to share structural similarities to previous ones. Another reason may be that scientific results are limited by the environments that we predominantly explore, while less focus thus-far having been paid to, for example, cyanobacteria from terrestrial or extreme environments. Compared to discoveries of natural products from other bacteria that started in the 1940s,^12^ cyanobacterial metabolite discoveries began only in the 1970s. The overall trend is especially driven by discoveries of peptide-based metabolites and common cyanopeptide classes make up more than one third of all peptidic metabolites in CyanoMetDB (Figure 2B). Within each class, compounds show high structural similarity supporting the previous observations that the contribution of novel structural scaffolds is typically low for natural products within the pool of cyanobacterial metabolites.^12^

### Chemical space

CyanoMetDB show that cyanobacterial metabolites occupy a wide range of molecular weights between 100 and 2500 Da. Of these, 57% are cyclic compounds and 65% are peptides. The mass range of known compounds does not perfectly fit a normal distribution (Figure 3A). Analysing peptidic metabolites separately demonstrates that peptides mainly account for the compounds with high molecular weights (Figure 3B). Note here that by peptidic, we refer strictly to the presence of at least one peptide bond, not the biosynthetic pathway.

**Figure 3.**
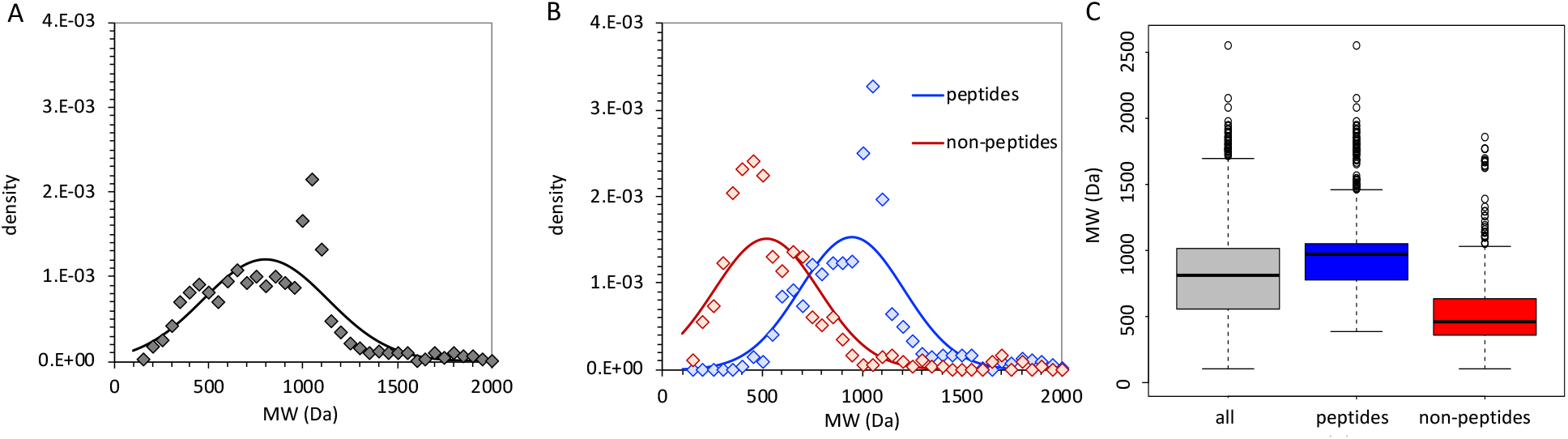
Probability density function of molecular weights for A) all compounds in the database (black), and; B) those containing a peptide bond (blue) and all other non-peptide-containing compounds (red), showing the distribution of the data (50-Da bins) as diamonds and the fitted normal distribution density function as solid lines. C) Box plot of molecular weights.

Among the peptides, more compounds are present in the 1000–1100 Da range than expected from a normally-distributed dataset. In particular, cyanopeptides can be classified to a large extent into common peptide classes including microcystins, anabaenopeptins, cyanopeptolins, and other depsipeptides that cover the majority of compounds with molecular weights above 900 Da (Figure 4). The distribution based on metabolite classes shows that microcystins and cyanopeptolins, in particular, contribute to the high abundance of known metabolites between 1000–1100 Da. Most likely this is because these classes have received more attention in studies focusing on identifying new members of these prominent classes that are often identified in the same species, or perhaps because microcystins and cyanopeptolins are especially abundant secondary metabolites from cyanobacteria.

**Figure 4.**
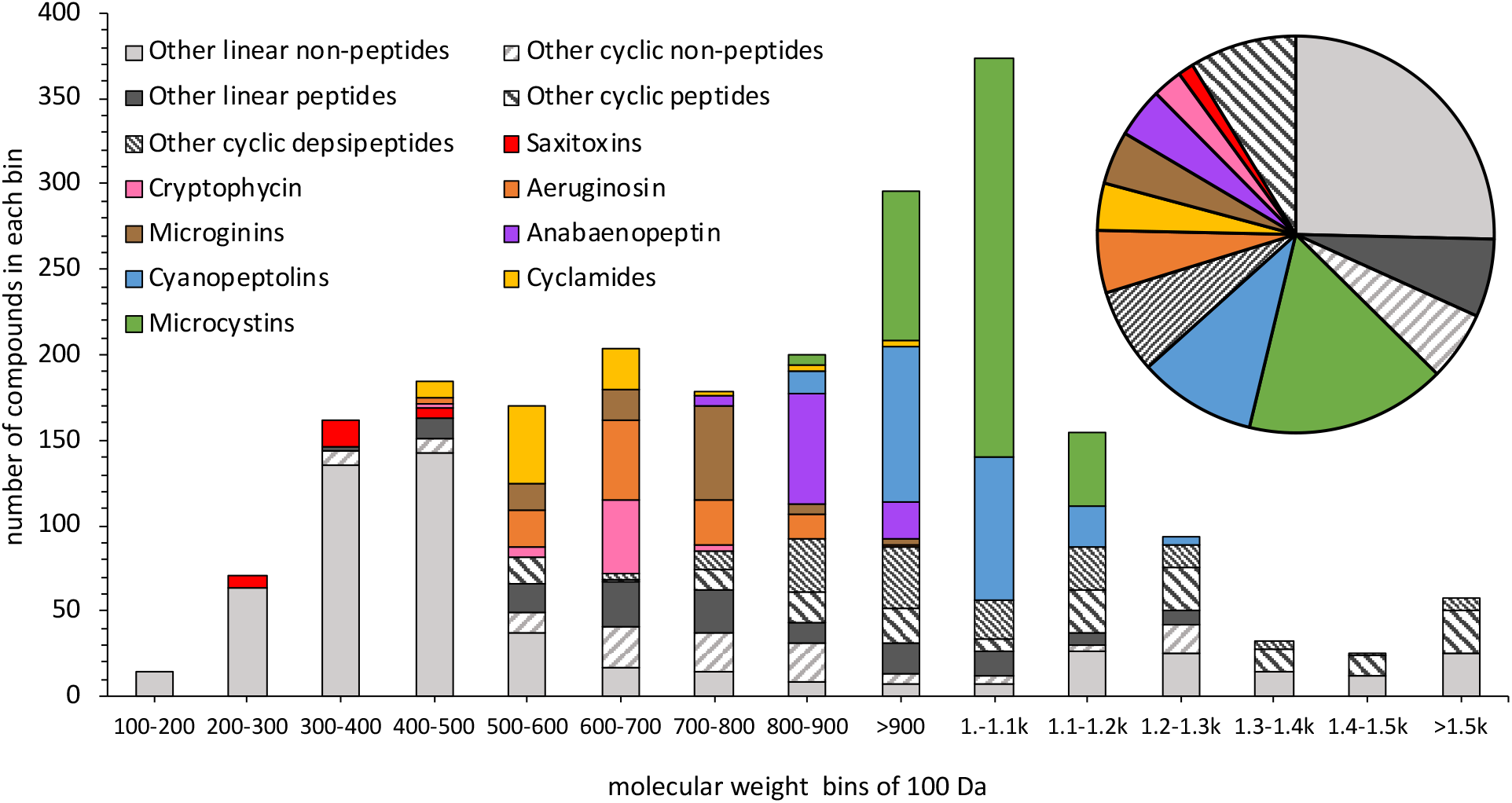
Distribution of molecular weights for secondary metabolites in the database from 100 to >1500 Da with 100-Da bin size showing the contribution of prominent metabolite classes and other linear and cyclic peptides and non-peptides, and the relative contribution to all database entries in the pie chart.

For the other non-peptide metabolites, we can see a particularly high contribution between 350–500 Da (Figure 3B). The molecular weight distribution of marine natural products has previously been shown to also center around 350 Da.^13^ In general, the non-peptide-based metabolites from cyanobacteria are more difficult to classify because they lack unique structural features. The structural information (SMILES codes) in the database allow for substructure searching to identify common molecular motifs. For example, of the non-peptide compounds, 15% contain an ester bond and 11% a proline ring. Most compounds show unsaturation with 64% carrying at least one aromatic group, 30% having 1–2 aromatic rings, and 6% having 3–6 aromatic rings. Halogen atoms are present in 30% of the non-peptide compounds (26% Cl, and 4% Br) and 20% contain sulfur. The structural information also allows calculations of other physico-chemical properties such as hydrogen bonding, aromaticity, or pKa values. However, for those calculations the applicability domain of any underlying models needs to be considered.

Secondary metabolites from cyanobacteria have mostly been discovered by “top-down” approaches from extraction of biomass and analyses that were guided by chemical motifs (peptides, molecular weight group, common fragments) or non-targeted searches by MS. Within the CyanoMetDB, 12% of compounds were first identified from field samples and the remaining compounds were identified from laboratory-grown cultures. In total, more than 50 different cyanobacterial genera were used, dominated by *Microcystis* (17% of all entries), *Moorea/Lyngbya*, *Nostoc*, *Anabaena/Dolichospermum*, *Oscillatoria/Planktothrix*, *Nodularia*, *Scytonema*, *Fischerella*, and *Symploca*, in decreasing order of total entries. These genera are not necessarily the main producers of the respective metabolites but were predominantly used for initial structure elucidation.

### Implications for suspect screening, biosynthetic and bioactivity-guided analysis

The CyanoMetDB dataset contains structurally known cyanobacterial metabolites that will aid future research including, but not limited to, suspect screening by MS, analysis of biosynthetic variation, bioactivity-guided analysis, and environmental behaviour.

#### Suspect Screening

Mass spectrometry-based analysis methods offer considerable opportunities and are the state-of-the-art analytical techniques to identify and quantify structurally-known compounds, i.e., targets and suspects. Targeted LC-MS analysis typically uses tandem mass spectrometry (MS/MS) and identifies compounds by characteristic fragment ions using the selected reaction monitoring scan mode of a low-resolution triple quadrupole mass spectrometer. Quantification by targeted analysis relies upon the knowledge of the characteristic fragments ions that are obtained from reference spectra or availability reference standards. Reference spectra and reference standards are, however, currently only available for a small fraction of entries in the CyanoMetDB (e.g., few microcystins, anatoxin-a, cylindrospermopsin, saxitoxins). Accordingly, targeted analyses are rather limited to assess the exact concentration in environmental samples for a wider range of cyanobacterial metabolite. Alternatively, LC-HRMS can be used for suspect screening of any compound with known molecular formula. One approach is by triggering fragmentation during MS/MS analysis of those spectral features in a sample that match the molecular formulae from a provided suspect list (i.e., an inclusion list). The fragments of a suspect can be matched against reference spectra or *in-silico* predictions to increase the level of confidence for the identification of a compound.^14^ With advances in machine learning and network-based analyses, further opportunities are also arising for the discovery of novel, or structurally-related, cyanobacterial metabolites based on holistic analysis and clustering of LC-HRMS/MS fragmentation datasets. These procedures all require a collated knowledgebase of structurally known compounds, which CyanoMetDB provides.

#### Biosynthetic basis

Discovery of secondary metabolites based-on genetic information is a promising “bottom up” approach that can reveal additional compounds,^15^ though this remains largely theoretical at this stage. Recent advances in microbial genomics have greatly improved our understanding of the biochemical mechanisms responsible for the biosynthesis of cyanobacterial natural products.^16^ The majority of these natural products are synthesized through secondary metabolic pathways that catalyze the synthesis of the complex secondary metabolites in coordinated enzyme cascades.^16^ Approximately 75% of the natural products included in the CyanoMetDB dataset can be assigned to structural families for which biosynthetic pathways have been reported from one or more representative. Many families of cyanobacterial secondary metabolites included in the CyanoMetDB exhibit extensive chemical variation but share a structural core that defines the family (Figure 1). Typically the biosynthesis of compounds in one family share a set of conserved biosynthetic enzymes for the synthesis of the defining structural core, but also a set of accessory tailoring enzymes that are not universally conserved.^16, 17^ The biosynthetic logic underlying the synthesis of most common cyanobacterial toxins, including microcystins,^18^ nodularins,^19^ saxitoxins,^20^ cylindrospermopsins,^21^ anatoxins,^22^ are now well understood. However, the genetic basis for the biosynthesis of specific secondary metabolite chemical variants is incomplete in many cases. Numerous molecular ecology methods have been developed to characterize the types of toxin producers in blooms based on the biosynthetic gene clusters.^17^ Compilation of CyanoMetDB has shown that the chemical variation of the major secondary metabolites is more extensive than previously realized. A more complete understanding of the biosynthetic basis for the chemical variation of secondary metabolites from cyanobacteria is necessary to ensure the unbiased detection of their biosynthetic pathways directly from environmental samples.

#### Bioactivity-guided analysis

With the fast-growing number of new secondary metabolites from cyanobacteria and the increasing awareness of their biological activities, a need for a comprehensive database is essential. Cyanobacterial metabolites present cytotoxic, antimicrobial, antifungal, antiprotozoal, enzyme inhibiting, anti-inflammatory, dermatotoxic and neurotoxic activities that can also be exploited by the pharmaceutical industry to develop new drugs potentially beneficial to humans.^24–26^ Future discoveries are likely to share structural similarities to previously discovered metabolites that may, however, exhibit differing potency and thus are critical to identify. Any modifications to the structure, such as replacement of amino acids by other residues, substitution by methylation, halogenation or oxidation and changes in configuration, can significantly affect the ability of cyanobacterial metabolites to evoke a biological response. For example, for the toxicity of microcystins, the cyclic structure and the Adda–D-Glu region, which interacts with the catalytic unit of protein phosphatases, is crucial. The modification of the less conserved positions 2 and 4 also plays a significant role for microcystin bioactivity (Figure 1).^6, 27^ The selective activity of cyanopeptolins against serine proteases depends on the residue in the Ahp-neighboring position. The Arg–Ahp-containing cyanopeptolin variants are mainly active against trypsin, whereas cyanopeptolins with hydrophobic amino acids (e.g., Phe, Tyr, Ile) show potent activity against toward chymotrypsin.^28^ In case of cryptophycins, which are cyclic depsipeptides composed of four fragments (i.e., A, B, C,, D), their effect on microtubule dynamics is determined by the intact 16-membered macrolide structure, reactive epoxide ring in unit A, methyl group in units A and C, *O*-methyl group and chloro-substituent in unit B, and isobutyl group in unit D.^29^ The CyanoMetDB presents a dataset including structural information for each metabolite that is deposited in a format immediately accessible to software tools (e.g., SMILES codes). The *in silico* studies, termed virtual screening (target-based or ligand-based), use such databases to overcome problems related to the limited availability of reference materials (i.e., pure compounds) and lack of information on their physico-chemical properties.^27^ In the case of new drug development from cyanopeptides, virtual screening will also help to reduce the cost and increase the success rate of the process. Chemoinformatics can also be helpful to discover as-yet unknown molecular targets of cyanobacterial metabolites in so-called target-fishing.^28,29^ The deposited structures in CyanoMetDB can be used as templates in these approaches and to design new chemical entities with desirable traits and ligand–target interactions.

#### Environmental behavior

As hundreds of secondary metabolites have been identified from cyanobacteria, we now face the questions: How toxic and abundant are the other cyanobacterial metabolites relative to known toxins? How stable are they in surface waters? How do their concentrations change during bloom events? Can they reach water treatment plant intakes? Answers to these questions are needed to quantify the exposure side of the risk equation and to prioritize cyanobacterial metabolites for toxicity testing, monitoring of surface waters, and evaluating removal in engineered water treatment systems. The inclusion of SMILES codes in CyanoMetDB offers the possibility of visualizing the planar or stereo-structure of a given compound. Access to the chemical structures enables use of chemical models to predict physico-chemical properties from quantitative structure–activity relationships. The structural information in CyanoMetDB also allows substructure searching for moieties with known reactivity, including biotic and abiotic processes. Open access tools that use known transformation rules to propose degradation of organic molecules can be applied to cyanobacterial metabolites of known structure as well. The predicted environmental transformation products can then also be included in virtual screening of bioactivity and suspect screening by MS as discussed above.

## Conclusions

Cyanobacterial metabolites have been studied for over 50 years, with the number of new discoveries continuing each year. In this work, the disparate data sources and primary research articles dealing with cyanobacterial metabolites have been collated and synthesized in to a freely-available and accessible flat-file database termed CyanoMetDB. The database comprises over 2000 entries, each corresponding to a unique cyanobacterial metabolite and its associated descriptors: name; molecular formula; molecular weight; monoisotopic mass; SMILES code; etc. CyanoMetDB represents a complementary tool to aid dereplication and analyses of cyanobacterial metabolites. We aim to continue this work as a community-driven effort in the future. As a next step, the content of this database will be integrated into other open access platforms to make it widely available and to benefit from already existing infrastructures and online tools. The database may also serve as a framework for connecting and collating other disparate data sources associated with cyanobacterial metabolites. This in turn may help to enrich collaborative research efforts, enhance the frequency with which compound annotations are assigned and aid communication, comparison, and interpretation of results.

## Supporting information

Database file CyanoMetDB_V01

Database file CyanoMetDB_References_V01

Database file CyanoMetDB_Microcystins_V01

## Acknowledgments

We thank São Paulo State Research Foundation (FAPESP - Grant No. 2014/50420-9) to E.P., University of São Paulo Foundation (FUSP-Grant No. 1979) to E.P.; the Coordination for the Improvement of Higher Education Personnel (CAPES - Grant No. 23038.001401/2018-92) to E.P.; CNPq (Grant No. 311048/2016-1 and 439065/2018-6) to E.P.; Marie Curie Innovative Training Network “Natural Toxins and Drinking Water Quality—From Source to Tap (NaToxAq, Grant No. 722493) funded by the European Commission to E.M.-L.J.; discretionary fund by Eawag to E.M.-L.J.; National Science Centre in Poland (Grant No. 2017/25/B/NZ9/00202 and 2019/33/B/NZ9/02018) to H.M.-M.; Novo Nordisk Foundation (18OC0034838) to D.P.F.; and NordForsk NCoE program “NordAqua” (Project No. 82845)) and Jane and Aatos Erkko Foundation to K.S.. We thank Roger Linginton and Jeffrey van Santen for extracting and supplying cyanobacterial metabolite entries from the Natural Product Atlas.

## References

1. Ibelings, B. W.; Backer, L. C.; Kardinaal, W. E. A.; Chorus, I., Current approaches to cyanotoxin risk assessment and risk management around the globe. Harmful Algae 2014, 40, 63–74.

2. Guidelines for Drinking Water Quality. World Health Organization: Geneva, 2004; Vol. Vol 1.

3. Le Manach, S.; Duval, C.; Marie, A.; Djediat, C.; Catherine, A.; Edery, M.; Bernard, C.; Marie, B., Global metabolomic characterizations of *Microcystis spp*. Highlights clonal diversity in natural bloom-forming populations and expands metabolite structural diversity. Front Microbiol 2019, 10.

4. van Santen, J. A.; Jacob, G.; Singh, A. L.; Aniebok, V.; Balunas, M. J.; Bunsko, D.; Neto, F. C.; Castano-Espriu, L.; Chang, C.; Clark, T. N.; Little, J. L. C.; Delgadillo, D. A.; Dorrestein, P. C.; Duncan, K. R.; Egan, J. M.; Galey, M. M.; Haeckl, F. P. J.; Hua, A.; Hughes, A. H.; Iskakova, D.; Khadilkar, A.; Lee, J. H.; Lee, S.; LeGrow, N.; Liu, D. Y.; Macho, J. M.; McCaughey, C. S.; Medema, M. H.; Neupane, R. P.; O’Donnell, T. J.; Paula, J. S.; Sanchez, L. M.; Shaikh, A. F.; Soldatou, S.; Terlouw, B. R.; Tran, T. A.; Valentine, M.; van der Hooft, J. J. J.; Vo, D. A.; Wang, M. X.; Wilson, D.; Zink, K. E.; Linington, R. G., The natural products atlas: an open access knowledge base for microbial natural products discovery. ACS Central Sci 2019, 5, (11), 1824–1833.

5. Meriluoto, J.; Spoof, L.; Codd, G. A., Handbook of Cyanobacterial Monitoring and Cyanotoxin Analysis. Wiley: 2017.

6. Bouaicha, N.; Miles, C. O.; Beach, D. G.; Labidi, Z.; Djabri, A.; Benayache, N. Y.; Nguyen-Quang, T., Structural diversity, characterization and toxicology of microcystins. Toxins 2019, 11, (12).

7. Miles, C. O.; Stirling, D., Toxin Mass List, version 16. Available online: https://www.researchgate.net/publication/337258461_Toxin_mass_list_COM_v160_microcystin_and_nodularin_lists_and_mass_calculators_for_mass_spectrometry_of_microcystins_nodularins_saxitoxins_and_anatoxins. Assessed on 14 April 2020. 2019.

8. Heller, S. R.; McNaught, A.; Pletnev, I.; Stein, S.; Tchekhovskoi, D., InChI, the IUPAC international chemical identifier. J Cheminformatics 2015, 7.

9. Crawford, J. M.; Mahlstedt, S. A.; Malcolmson, S. J.; Clardy, J.; Walsh, C. T., Dihydrophenylalanine: a prephenate-derived photorhabdus luminescens antibiotic and intermediate in dihydrostilbene biosynthesis. Chem Biol 2011, 18, (9), 1102–1112.

10. Parker, J. B.; Walsh, C. T., Stereochemical outcome at four stereogenic centers during conversion of prephenate to tetrahydrotyrosine by BacABGF in the bacilysin pathway. Biochemistry 2012, 51, (28), 5622–5632.

11. Welker, M.; von Döhren, H., Cyanobacterial peptides - Nature’s own combinatorial biosynthesis. FEMS Microbiol Rev 2006, 30, (4), 530–563.

12. Pye, C. R.; Bertin, M. J.; Lokey, R. S.; Gerwick, W. H.; Linington, R. G., Retrospective analysis of natural products provides insights for future discovery trends. Proc Natl Acad Sci USA 2017, 114, (22), 5601–5606.

13. Blunt, J. W.; Copp, B. R.; Keyzers, R. A.; Munro, M. H. G.; Prinsep, M. R., Marine natural products. Nat Prod Rep 2017, 34, (3), 235–294.

14. Schymanski, E. L.; Jeon, J.; Gulde, R.; Fenner, K.; Ruff, M.; Singer, H. P.; Hollender, J., Identifying small molecules via high resolution mass spectrometry: communicating confidence. Environ Sci Technol 2014, 48, (4), 2097–2098.

15. Moosmann, P.; Ueoka, R.; Gugger, M.; Piel, J., Aranazoles: extensively chlorinated nonribosomal peptide-polyketide hybrids from the cyanobacterium *Fischerella* sp. PCC 9339. Org. Lett. 2018, 20, (17), 5238–5241.

16. Dittmann, E.; Gugger, M.; Sivonen, K.; Fewer, D. P., Natural product biosynthetic diversity and comparative genomics of the cyanobacteria. Trends Microbiol 2015, 23, (10), 642–652.

17. Dittmann, E.; Fewer, D. P.; Neilan, B. A., Cyanobacterial toxins: biosynthetic routes and evolutionary roots. FEMS Microbiol Rev 2013, 37, (1), 23–43.

18. Tillett, D.; Dittmann, E.; Erhard, M.; von Döhren, H.; Borner, T.; Neilan, B. A., Structural organization of microcystin biosynthesis in *Microcystis aeruginosa* PCC7806: an integrated peptide-polyketide synthetase system. Chem Biol 2000, 7, (10), 753–764.

19. Moffitt, M. C.; Neilan, B. A., Characterization of the nodularin synthetase gene cluster and proposed theory of the evolution of cyanobacterial hepatotoxins. Appl Environ Microb 2004, 70, (11), 6353–6362.

20. Kellmann, R.; Mihali, T. K.; Neilan, B. A., Identification of a saxitoxin biosynthesis gene with a history of frequent horizontal gene transfers. J Mol Evol 2009, 68, (3), 292–292.

21. Mihali, T. K.; Kellmann, R.; Muenchhoff, J.; Barrow, K. D.; Neilan, B. A., Characterization of the gene cluster responsible for cylindrospermopsin biosynthesis. Appl Environ Microb 2008, 74, (3), 716–722.

22. Mejean, A.; Mann, S.; Maldiney, T.; Vassiliadis, G.; Lequin, O.; Ploux, O., Evidence that biosynthesis of the neurotoxic alkaloids anatoxin-a and homoanatoxin-a in the cyanobacterium *Oscillatoria* PCC 6506 occurs on a modular polyketide synthase initiated by L-proline. J Am Chem Soc 2009, 131, (22), 7512–+.

23. Demay, J.; Bernard, C.; Reinhardt, A.; Marie, B., Natural products from cyanobacteria: focus on beneficial activities. Mar Drugs 2019, 17, (6).

24. Singh, R. K.; Tiwari, S. P.; Rai, A. K.; Mohapatra, T. M., Cyanobacteria: an emerging source for drug discovery. J Antibiot 2011, 64, (6), 401–412.

25. S., K.; Divyashree, M.; Mani, M.; Mamatha, B., Algae and cyanobacteria as a source of novel bioactive compounds for biomedical applications. In Advances in Cyanobacterial Biology, Academic Press, 2020; pp 173–194.

26. Huang, I. S.; Zimba, P. V., Cyanobacterial bioactive metabolites – a review of their chemistry and biology (vol 83, pg 42, 2019). Harmful Algae 2019, 86, 138–138.

27. Fontanillo, M.; Kohn, M., Microcystins: synthesis and structure-activity relationship studies toward PP1 and PP2A. Bioorg Med Chem 2018, 26, (6), 1118–1126.

28. Yamaki, H.; Sitachitta, N.; Sano, T.; Kaya, K., Two new chymotrypsin inhibitors isolated from the cyanobacterium *Microcystis aeruginosa* NIES-88. J Nat Prod 2005, 68, (1), 14–18.

29. Golakoti, T.; Ogino, J.; Heltzel, C. E.; LeHusebo, T.; Jensen, C. M.; Larsen, L. K.; Patterson, G. M. L.; Moore, R. E.; Mooberry, S. L.; Corbett, T. H.; Valeriote, F. A., Structure determination, conformational analysis, chemical stability studies, and antitumor evaluation of the cryptophycins. Isolation of 18 new analogs from *Nostoc sp* strain CSV 224 (vol 117, pg 12030, 1995). J Am Chem Soc 1996, 118, (13), 3323–3323.

